# Fcγ receptor-dependent antibody effector functions are required for vaccine protection against infection by antigenic variants of SARS-CoV-2

**DOI:** 10.1101/2022.11.27.518117

**Authors:** Samantha R. Mackin, Pritesh Desai, Bradley M. Whitener, Courtney E. Karl, Meizi Liu, Ralph S. Baric, Darin K. Edwards, Taras M. Chicz, Ryan P. McNamara, Galit Alter, Michael S. Diamond

## Abstract

Emerging SARS-CoV-2 variants with antigenic changes in the spike protein are neutralized less efficiently by serum antibodies elicited by legacy vaccines against the ancestral Wuhan-1 virus. Nonetheless, these vaccines, including mRNA-1273 and BNT162b2, retained their ability to protect against severe disease and death, suggesting that other aspects of immunity control infection in the lung. Although vaccine-elicited antibodies can bind Fc gamma receptors (FcγRs) and mediate effector functions against SARS-CoV-2 variants, and this property correlates with improved clinical COVID-19 outcome, a causal relationship between Fc effector functions and vaccine-mediated protection against infection has not been established. Here, using passive and active immunization approaches in wild-type and Fc-gamma receptor (FcγR) KO mice, we determined the requirement for Fc effector functions to protect against SARS-CoV-2 infection. The antiviral activity of passively transferred immune serum was lost against multiple SARS-CoV-2 strains in mice lacking expression of activating FcγRs, especially murine FcγR III (CD16), or depleted of alveolar macrophages. After immunization with the preclinical mRNA-1273 vaccine, protection against Omicron BA.5 infection in the respiratory tract also was lost in mice lacking FcγR III. Our passive and active immunization studies in mice suggest that Fc-FcγR engagement and alveolar macrophages are required for vaccine-induced antibody-mediated protection against infection by antigenically changed SARS-CoV-2 variants, including Omicron strains.

## INTRODUCTION

Since the emergence of severe acute respiratory syndrome coronavirus 2 (SARS-CoV-2) in late 2019, 638 million infections and 6.6 million deaths have been reported (https://covid19.who.int/). As part of the global response to COVID-19, vaccines using multiple different platforms (mRNA, adenoviral-vectored, subunit-based, and inactivated virion) were generated and deployed resulting in reductions in symptomatic infections, hospitalizations, and deaths. These SARS-CoV-2 vaccines all targeted the viral spike protein derived from strains that circulated in early 2020. However, the continuing evolution of SARS-CoV-2, with increasing numbers of amino acid changes in the spike protein amidst successive waves of infection, has jeopardized the immunity generated by these vaccines and the control of virus infection and transmission^1^.

SARS-CoV-2 vaccination can induce neutralizing antibodies that inhibit infection^2,3^. Correlates of vaccine protection initially focused on the neutralizing activity of elicited anti-spike antibodies^4,5^. Emerging variants of concern, which have amino acid substitutions in regions targeted by neutralizing antibodies including the receptor binding domain (RBD) and N terminal domain (NTD)^6,7^ have jeopardized vaccine-mediated protection against infection and prompted the development of bivalent vaccine boosters^8^. Indeed, a substantial decrease in the neutralizing activity of serum antibodies elicited by vaccines against the ancestral Wuhan-1 virus has been observed against emerging variants, which has correlated with symptomatic breakthrough infections, especially with Omicron lineage viruses^9–11^. The large number (>30) of substitutions in the spike protein in Omicron lineage strains, which abrogates or reduces binding of the majority of highly neutralizing vaccine-derived and therapeutic antibodies, has been termed an antigenic shift^12–14^. Despite the loss in serum neutralizing activity against variants such as those in Omicron lineage, most individuals remain protected against severe disease and death. The basis for this protection has not been fully determined but could be due to beneficial effects of non-neutralizing antibodies, cross-reactive T cell responses, or anamnestic memory B cell responses^6,15–18^.

Fc effector function activity of non-neutralizing, cross-reactive, anti-spike antibodies is one hypothesized mechanism for protection against antigenically-shifted SARS-CoV-2 variants^7^. In patients with moderate to severe COVID-19, the ability of antibodies to bind Fc-gamma receptors (FcγR) and mediate effector functions correlated with increased survival^19^. Interactions of the conserved Fc region of IgG antibodies with FcγR or complement can promote clearance of virally-infected cells through antibody-dependent cellular cytotoxicity (ADCC), antibody-dependent cellular phagocytosis (ADCP), or complement-dependent deposition and phagocytosis or lysis. Indeed, monoclonal antibodies (mAbs) that lose their ability to neutralize SARS-CoV-2 variants yet still bind the spike protein avidly enough to trigger Fc effector functions retain protective activity^15,20^. Analogously, non-neutralizing antibodies induced by SARS-CoV-2 vaccines have been linked to protection against variant Omicron strains by virtue of their ability to engage FcγRs and promote clearance^21–23^. Furthermore, the depletion of RBD-specific antibodies from serum of mRNA-1273 or BNT162b2 vaccinated individuals did not appreciably impact Fc-mediated effector function activity in cell culture^6^, suggesting that antibodies recognizing conserved, non-neutralizing epitopes may contribute to protection against variant strains.

Although serum-derived anti-SARS-CoV-2 antibody-mediated Fc effector functions can activate complement deposition, immune cell phagocytosis, and target cell killing *in vitro*, their contribution to protection *in vivo* remains uncertain. Existing data on the role of Fc-FcγR interactions in the context of vaccine-mediated protection against SARS-CoV-2 is largely correlative. To address this gap, we evaluated the impact of Fc effector functions in the context of passive transfer of vaccine-elicited antibodies or active immunization with mRNA-1273 vaccine using wild-type, C1q KO, and FcγR KO C57BL/6 mice and challenge with SARS-CoV-2 viruses. In passive serum transfer experiments, we found that activating FcγRs but not C1q were required to control SARS-CoV-2 infection in the lower respiratory tract, and protection was lost in mice depleted of alveolar macrophages but not neutrophils and monocytes. Experiments with mice lacking individual FcγRs showed the protective effect of passively transferred serum antibody on viral load reduction required expression of FcγR III. To determine the impact of FcγRs in the context of active immunization, wild-type, FcγR I KO, FcγR III KO, and FcγR I/III/IV KO mice were administered a two-dose primary vaccination series with mRNA-1273, evaluated for immunogenicity, and then challenged with the antigenically shifted Omicron BA.5 strain. Although the levels of anti-RBD antibody, neutralizing antibody, and spike-specific T cells were similar in all tested strains of mice, protection against infection in the nasal turbinates and lungs was lost in FcγR III KO and FcγR I/III/IV KO mice. Overall, our results in mice suggest that Fc-FcγR interactions contribute to antiviral protection *in vivo* in the context of both passive and active immunization with legacy vaccines, particularly when neutralizing antibody levels are low against antigenically distant SARS-CoV-2 variant strains.

## RESULTS

### Immunoglobulin subclass and FcγR binding of vaccine-induced immune serum

To begin to evaluate the contribution of Fc effector functions to antibody protection against SARS-CoV-2 infection, we profiled vaccine-induced antibodies from sera pooled from immunized C57BL/6 mice using a systems serology assay^24^. We measured the binding of polyclonal antibodies to several spike proteins (Wuhan-1, B. 1.617.2, BA.1, and BA.4/5) and determined their IgG subclass specificity (IgG1, IgG2a, or IgG2c) and ability to interact with specific FcγRs (FcγR IIb, FcγR III, or FcγR IV). We used naïve sera and binding to influenza hemagglutinin (HA) protein as negative controls (**Fig 1a-f**). Immune sera contained higher levels of IgG1, IgG2b, and IgG2c antibodies against Wuhan-1, B.1.617.2, BA.1, and BA.4/.5 but not HA compared to naïve sera (**Fig 1a-c**). Anti-spike antibody binding to inhibitory (FcγR IIb) and activating (FcγR III and FcγR IV) was higher in immune sera compared to naïve sera for all SARS-CoV-2 spike variants (**Fig 1d-f**). We also assessed the effector function activity of immune serum using assays that measure spike-specific antibody-dependent cellular phagocytosis in murine monocytes (ADCP) and neutrophils (ADNP) (**Fig 1g-h and Extended Data Fig 1**). Compared to naïve sera, vaccine-elicited immune sera promoted greater ADCP and ADNP activity against the Wuhan-1 and BA.4/.5 spike proteins. In comparison, immune sera did not enhance antibody-dependent natural killer cell activation (CD107a expression, **Fig 1i**). Immune sera also promoted antibody-dependent complement deposition (ADCD) on beads coated with spike proteins compared to HA protein (**Fig 1j**). Overall, these studies indicated that our pooled vaccine sera have a diversity of antibodies against spike proteins that enables binding to FcγRs, and most Fc-mediated effector functions in cell culture.

**Figure 1.**
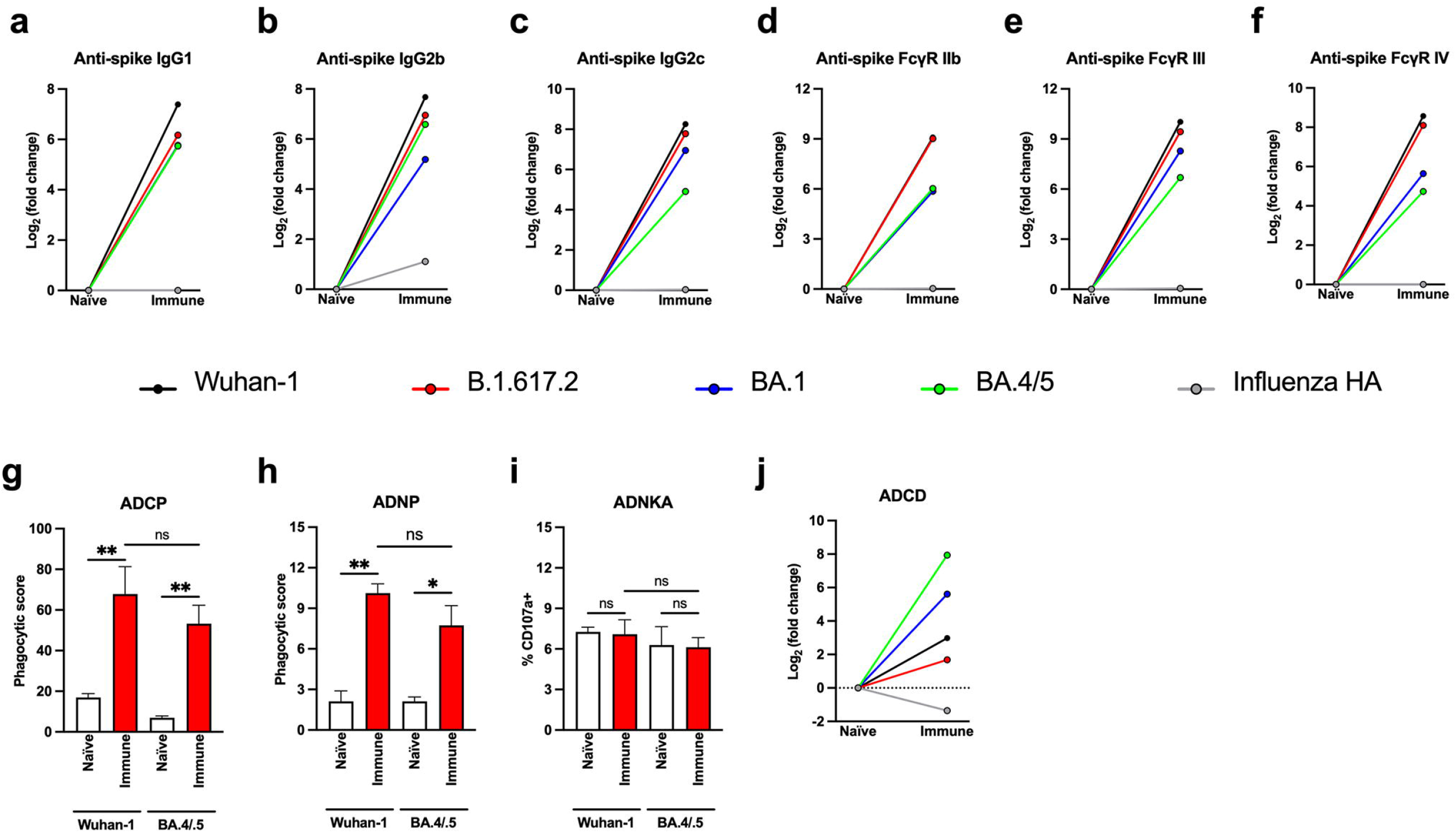
Systems serology analysis of vaccine-induced immune sera. (**a-c**) Levels of IgG1 (**a**), IgG2b (**b**), IgG2c (**c**) that bind to SARS-CoV-2 spike [Wuhan-1, B.1.617.2, BA.1, and BA.4/5], or influenza hemagglutinin (HA) in naïve and vaccine-induced immune sera. (**d-f**) Levels of spike- or HA-binding IgG antibodies that engage FcγR IIb (**d**), FcγR III (**e**), or FcγR IV (**f**) in naïve and vaccine-induced immune sera. (**g-j**) Antibody effector functions. Antibody-mediated cellular phagocytosis with monocytes (ADCP, **g**) or neutrophils (ADNP, **h**) activity using vaccine-induced immune (red) or naïve (white) sera and beads coated with SARS-CoV-2 Wuhan-1 and BA.4/5 spike proteins and murine monocytes (bars indicate median values; one-way ANOVA with Tukey’s post-test; ns, not significant; *\*P* < 0.05, \*\**P* < 0.01). (**i**) CD107a surface expression on natural killer cells (ADNKA) after incubation with beads encoded with Wuhan-1 or BA4/5 spike proteins and immune sera (bars indicate median values; one-way ANOVA with Tukey’s post-test; ns, not significant). (**j**) Deposition of complement (ADCD) on beads coated with indicated SARS-CoV-2 spike or influenza HA proteins after treatment with naïve or immune sera.

### Protection in the lungs conferred by passive sera transfer requires Fc-FcγR engagement

To assess the impact of Fc effector functions *in vivo* in the context of polyclonal immune anti-SARS-CoV-2 antibodies, pooled naïve or vaccine-elicited immune sera was transferred passively to 12-week-old male wild-type, FcγR I/III/IV KO (common γ chain KO), or C1q KO C57BL/6 mice; FcγR I/III/IV KO mice lack the common γ chain present in all activating murine FcγRs, whereas C1q KO mice lack C1q, a protein required for initiation of the antibody-dependent complour experiments, we focusedement activation pathway. One day after transfer, mice were inoculated with SARS-CoV-2 MA-10^25^, and 4 days post-infection (dpi), nasal wash, nasal turbinates, and lungs were harvested (**Fig 2a**). We used the mouse-adapted MA-10 strain for these initial studies because it spreads to the lungs of conventional C57BL/6 mice without a need for ectopic human ACE2 expression. Pooled vaccine-induced immune sera neutralized MA-10 at a 1/3,300 dilution titer, and one day after transfer, serum from recipient mice neutralized MA-10 with a titer of 1/36 (**Fig 2b**). In the nasal washes and nasal turbinates of the upper airway of wild-type, FcγR I/III/IV KO, and C1q KO mice, we observed no significant differences in levels of viral RNA or infectious virus among the three groups receiving naive or immune serum. Although there was a trend towards less viral infection in the nasal turbinates of animals receiving immune compared to naïve sera, the comparisons did not reach statistical significance (**Fig 2c-e**); these results are consistent with the lower accumulation of IgG in the upper respiratory tract after passive transfer by a systemic route^26^. Results from lung tissues, however, showed a different pattern, with a loss of protection against infection by immune sera (viral RNA levels and infectious virus) in FcγR I/III/IV KO but not in C1q KO mice (**Fig 2f-g**). Thus, FcγR expression in the lower respiratory tract appears important for control of infection after passive antibody transfer.

**Figure 2.**
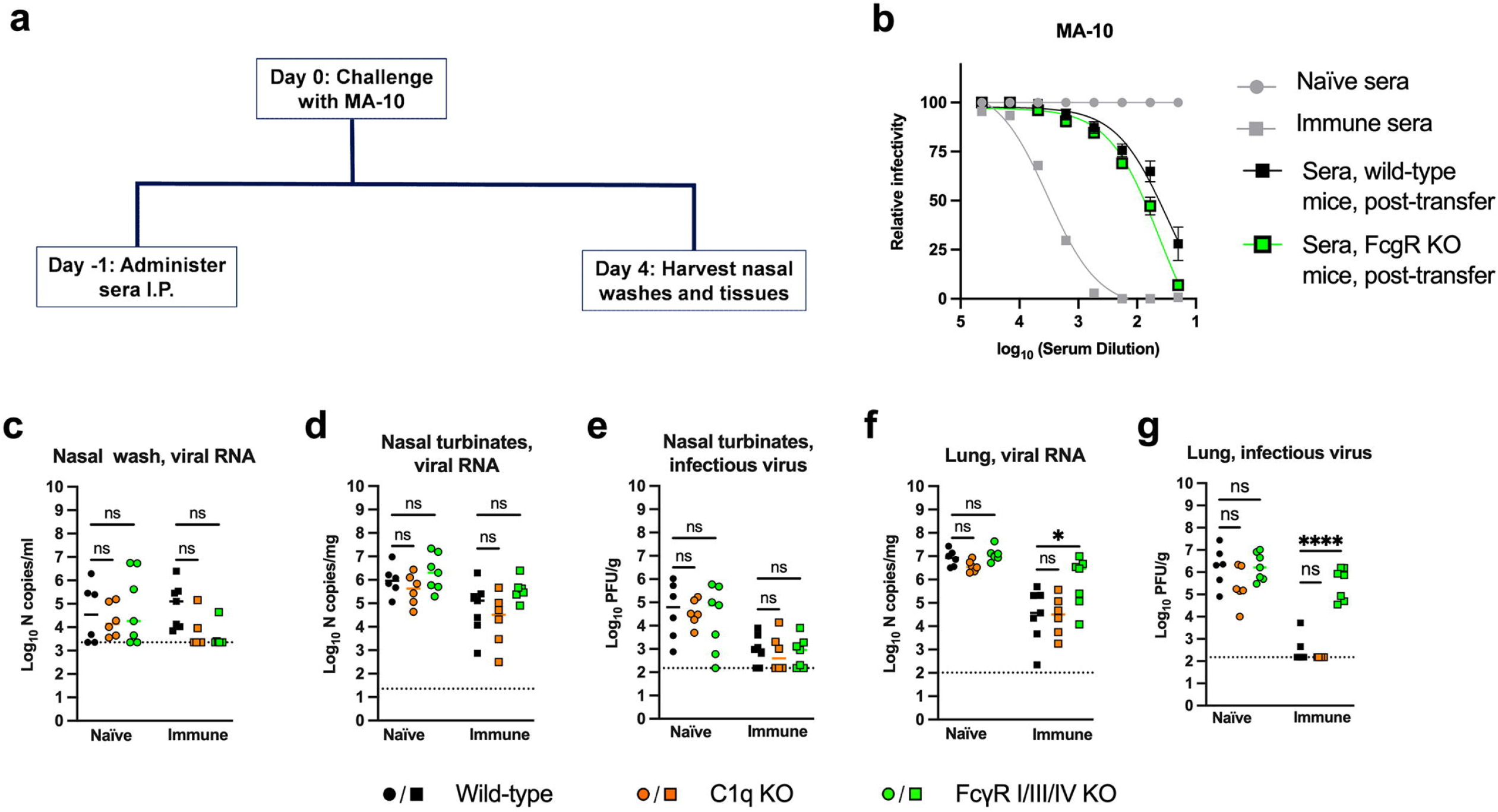
Vaccine-derived immune sera protection against SARS-CoV-2 MA-10 infection in wild-type C1q KO, and FcγR I/III/IV KO mice. (**a**) Scheme of passive transfer, virus challenge and tissue harvest. (**b**) Neutralizing antibody responses against SARS-CoV-2 MA-10 using sera from naïve (circles) or Wuhan-1 spike protein vaccinated mice (pooled from animals immunized and boosted with mRNA-1273 or ChAd-SARS-CoV-2-S) (squares). Also shown is serum neutralizing antibody activity from recipient wild-type (black squares) and FcγR I/III/IV KO (green squares) mice one day after transfer of immune sera. (**c-g**) Twelve-week-old male wild-type, C1q KO, and FcγR I/III/IV KO C57BL/6 mice were passively transferred by intraperitoneal injection 60 μL of naïve or vaccine-induced immune sera one day before intranasal challenge with 10^3^ FFU of SARS-CoV-2 MA-10. At 4 dpi, viral RNA in the nasal wash (**c**), nasal turbinates (**d**), and lungs (**f**) were quantified by qRT-PCR, and infectious virus in the nasal turbinates (**e**) and lungs (**g**) was determined by plaque assay (bars indicate median values; n = 6-7 mice per group, two experiments, dotted lines show limit of detection [LOD]). One-way ANOVA with Tukey’s post-test; ns, not significant; \**P* < 0.05, \*\*\*\**P* < 0.0001).

### Antibody protection in the lungs requires FcγR III engagement

Because MA-10 is not matched to the vaccine antigen, in the context of passive transfer, we might underestimate the protection against infection afforded by serum neutralizing antibody. To address this issue and also identify which FcγR contributed to the protective phenotype, we passively transferred naïve or immune sera to 12-week-old male wild-type, FcγR I KO, FcγR II KO, FcγR III KO, and FcγR I/III/IV KO congenic C57BL/6 male mice before inoculation with SARS-CoV-2 WA1/2020 N501Y/D614G (**Fig 3a**), a more closely matched virus; because this suite of FcγR-deficient C57BL/6 mice lacks human ACE2 expression, we used a virus with a mouse-adapting N501Y mutation^27,28^. Pooled vaccine-elicited immune sera neutralized WA1/2020 N501Y/D614G more efficiently than MA-10 at a 1/16,750 serum dilution, and one day after transfer, serum from recipient mice neutralized WA1/2020 N501Y/D614G with a titer of 1/750 (**Fig 3b**), which exceeds a ~1/50 presumptive correlate of protection in humans^4^. As seen with MA-10 infection, in the nasal washes and nasal turbinates of the upper respiratory tract, we did not observe serum antibody protection in wild-type C57BL/6 mice (**Fig 3c-e**); thus, we focused analysis on the lung. Indeed, passive transfer of immune sera protected against SARS-CoV-2 infection in the lungs of wild-type C57BL/6 mice as measured by viral RNA and infectious virus levels (**Fig 3f-g**). Similar levels of protection were observed in FcγR I KO and FcγR II KO. However, protection against SARS-CoV-2 lung infection was diminished or lost in FcγR III and FcγR I/III/IV KO mice. These data suggest that even in the context of passive transfer of immune sera with neutralizing activity, protection against lower respiratory tract infection by SARS-CoV-2 is mediated at least in part by Fc interactions with activating FcγRs, particularly FcγR III.

**Figure 3.**
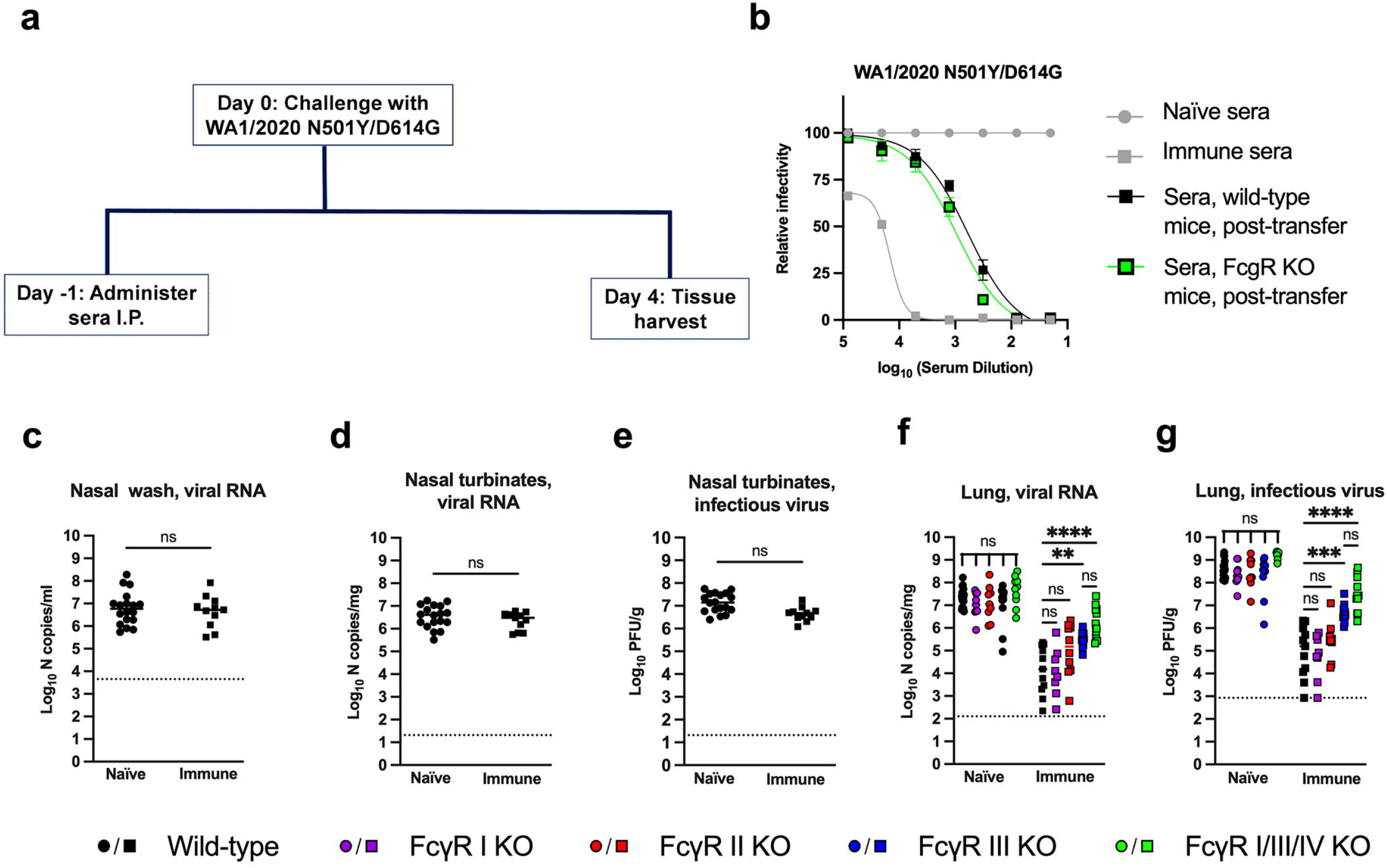
Vaccine-elicited immune sera protection against SARS-CoV-2 WA1/2020 N501/D614G infection in wild-type, FcγR I KO, FcγR II KO, FcγR III KO, and FcγR I/III/IV KO mice. (**a**) Scheme of passive transfer, virus challenge, and tissue harvest. (**b**) Neutralizing antibody response against SARS-CoV-2 WA1/2020 N501Y/D614G using sera from naïve (circles) or Wuhan-1 spike protein vaccinated (squares) mice. Also shown is serum neutralizing antibody activity from recipient wild-type (black squares) and FcγR I/III/IV KO (green squares) mice one day after transfer of immune sera. (**c-g**) Twelve-week-old male wild-type, FcγR I KO, FcγR II KO, FcγR III KO, and FcγR I/III/IV KO mice were passively transferred by intraperitoneal injection 60 μL of naïve or vaccine-immune sera one day before intranasal challenge with 10^4^ FFU of WA1/2020 N501Y/D614G. At 4 dpi, viral RNA and infectious virus was measured in the upper respiratory tract (nasal wash) (**c**), nasal turbinates (**d-e**)) or lungs (**f-g**) and quantified by qRT-PCR and plaque assay. Panels **c-e**: wild-type mice only; panels **f-g**: wild-type, FcγR I KO, FcγR II KO, FcγR III KO, and FcγR I/III/IV KO mice (bars indicate median; n = 9-18 mice per group, three experiments, dotted lines show LOD). One-way ANOVA with Tukey’s post-test (ns, not significant; \*\*\**P* < 0.01, \*\**P* < 0.001, \*\*\*\**P* < 0.0001).

### Vaccine-elicited immunity requires Fc-FcγR engagement to confer protection against SARS-CoV-2 infection

We next evaluated the dependence on Fc-FcγR engagement on protection against SARS-CoV-2 infection in the context of vaccine-elicited immunity, which induces both cellular and humoral responses. We immunized groups of nine-week-old male wild-type, FcγR I KO, FcγR III KO, and FcγR I/III/IV KO C57BL/6 mice twice over four weeks with 0.25 μg of control or preclinical mRNA-1273 vaccine (**Fig 4a**); we did not immunize FcγR II KO mice, since the virological phenotypes in the context of passive antibody transfer were present in mice lacking activating FcγRs but not FcγR II (**Fig 2 and 3**). The 0.25 μg dose of mRNA vaccine was used because the B and T cell responses generated in C57BL/6 mice with this dose approximate those observed in humans receiving 100 μg doses^29,30^. One potential limitation of this experiment is that a loss of activating FcγRs could affect vaccine-induced immune responses, which might confound interpretation of challenge studies. To evaluate this first, twenty-four days following boosting, serum was obtained to measure binding and neutralizing antibody against WA1/2020 N501Y/D614G and BA.5. As expected, higher levels of serum IgG were detected against Wuhan-1 than BA.5 receptor binding domain (RBD) protein (**Fig 4b-c**), consistent with the antigenic shift of Omicron lineage strains^12,14^. However, no statistical differences in binding titers were observed between the groups of vaccinated wildtype and FcγR KO mice (**Fig 4b-c**). Similarly, lower neutralization titers were detected against BA.5 than WA1/2020 N501Y/D614G, with no substantive differences observed between groups of vaccinated wild-type and FcγR KO (**Fig 4d-e**). Thus, mRNA-1273 vaccination induced relatively similar humoral immune responses in mice that were sufficient or deficient in FcγR expression. We also measured spike-specific CD4^+^ and CD8^+^ T cell responses in vaccinated mice using previously defined immunodominant peptides^31,32^. As expected, antigen-specific T cell responses were greater in animals give the mRNA-1273 vaccine compared to the control mRNA vaccine. However, IFN-γ and TNF-α responses in CD4^+^ and CD8^+^ T cells after peptide restimulation were equivalent in FcγR KO and wild-type mice after mRNA-1273 vaccination (**Extended Data Fig 2**). These experiments establish that FcγR KO and wild-type mice have similar serum antibody and T cell responses after mRNA vaccination.

**Figure 4.**
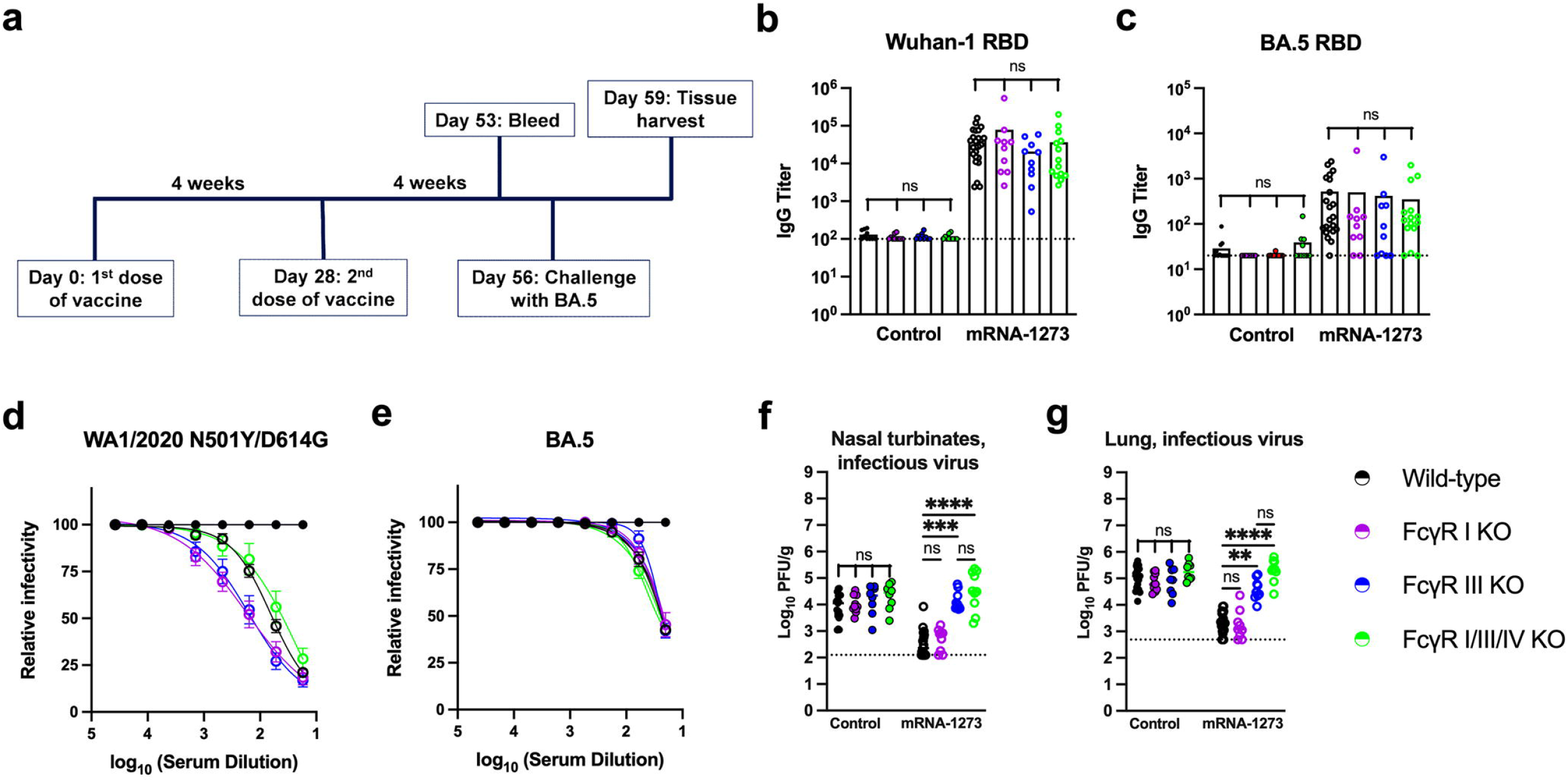
Protection against SARS-CoV-2 BA.5 infection after mRNA-1273 vaccination of wild-type and FcγR-deficient C57BL/6 mice. (**a**) Scheme of immunization, serum sampling, virus challenge, and tissue harvest. (**b-c**) Anti-Wuhan-1 (**b**) and BA.4/5 (**c**) RBD IgG responses from serum of mice immunized with control or mRNA-1273 vaccines (n = 10-22, boxes illustrate geometric mean titers [GMT], dotted lines show LOD). (**d-e**) Neutralizing antibody responses against WA1/2020 N501Y/D614G (**d**) and BA.5 (**e**) from serum collected from wild-type, FcγR I KO, FcγR III KO, and FcγR I/III/IV KO mice 25 days after completion of a two-dose primary vaccination series with control (closed circles) or mRNA-1273 (open circles). (**f-g**) Nine-week-old male wild-type, FcγR I KO, FcγR III KO, and FcγR I/III/IV KO mice were immunized twice at four-week intervals with control or mRNA-1273 vaccine via intramuscular route. Four weeks after the primary vaccination series, mice were challenged via intranasal route with 10^4^ FFU of BA.5. At 3 dpi, infectious virus in the nasal turbinates (**f**) and lungs (**g**) was determined by plaque assay (bars indicate median values; n = 8-10 mice per group, two experiments, dotted lines show LOD, one-way ANOVA with Dunnett’s test (ns, not significant, \*\**P* < 0.01, \*\*\**P* < 0.0001, \*\*\*\**P* < 0.0001).

To assess whether mice expressing FcγRs were differentially protected against SARS-CoV-2 infection by vaccine-induced immunity, animals were challenged by intranasal route with 10^3^ FFU of BA.5, and infectious virus in the nasal turbinates and lungs was measured at 3 dpi (**Fig 4a**). For these experiments, we used BA.5 as the challenge virus because: (i) it encodes a mouse-adapting mutation (N501Y) that facilitates replication in mice lacking human ACE2 expression^33^; (ii) it allowed us to assess protection against infection under conditions when high levels of neutralizing antibody are absent (**Fig 4e**); and (iii) BA.5, and other Omicron variants are circulating, so use of this strain could provide insight as to how legacy vaccines directed against ancestral spikes protect against severe BA.5 disease in humans. Notably, we observed protection against BA.5 infection in the upper and lower respiratory tract of wild-type and FcγR I KO mice but not in FcγR III KO or FcγR I/III/IV KO mice (**Fig 4f-g**). Thus, protection elicited by mRNA-1273 vaccine-induced immunity against the antigenically shifted BA.5 SARS-CoV-2 requires Fc-FcγR engagement, and FcγR III interactions in particular contribute to this phenotype in mice.

### Alveolar macrophages are required for antibody protection against BA.5 infection after passive immunization

We next addressed which FcγR III-expressing immune cells in the lung were important for mediating antibody protection in the context of passive transfer of immune sera and BA.5 challenge (**Fig 5**). Mice that received vaccine-elicited immune sera had detectable amounts of anti-BA.5 spike IgG but low levels of neutralizing activity (~1/10 titer), as expected (**Extended Data Fig 3**). Flow cytometric analysis of CD45^+^ immune cells in the lungs of wild-type C57BL/6 mice showed that murine monocytes, neutrophils, interstitial macrophages, and alveolar macrophages express FcγR III (**Extended Data Fig 4 and Supplementary Table 1**). We first assessed the role of neutrophils and monocytes by depleting these cells in wild-type mice with an anti-Ly6C/Ly6G (Gr-1) antibody (**Fig 5a**). Depletion of these cells in circulation (**Fig 5b-d, Extended Data Fig 5a**), which corresponds to depletion in the lung^34^, did not impact BA.5 infection in the nasal turbinates or lungs; decreased levels of infectious BA.5 virus in the lungs were seen after immune sera transfer regardless of whether neutrophils and monocytes were present (**Fig 5e-f**). We next depleted alveolar macrophages (**Fig 5g-i, Extended Data Fig 5b**) in the lung using a previously described protocol of intranasal administration of clodronate liposomes^35^. Treatment with clodronate, but not control liposomes, which depleted alveolar macrophages but not other FcγR-expressing immune cells in the lung, was associated with a loss of protection against BA.5 infection after passive transfer of immune but not non-immune (naïve) sera (**Fig 3j-k**). Together, these experiments establish an important role for FcγR III-expressing alveolar macrophages in antibody-mediated control of BA.5 infection in mice.

**Figure 5.**
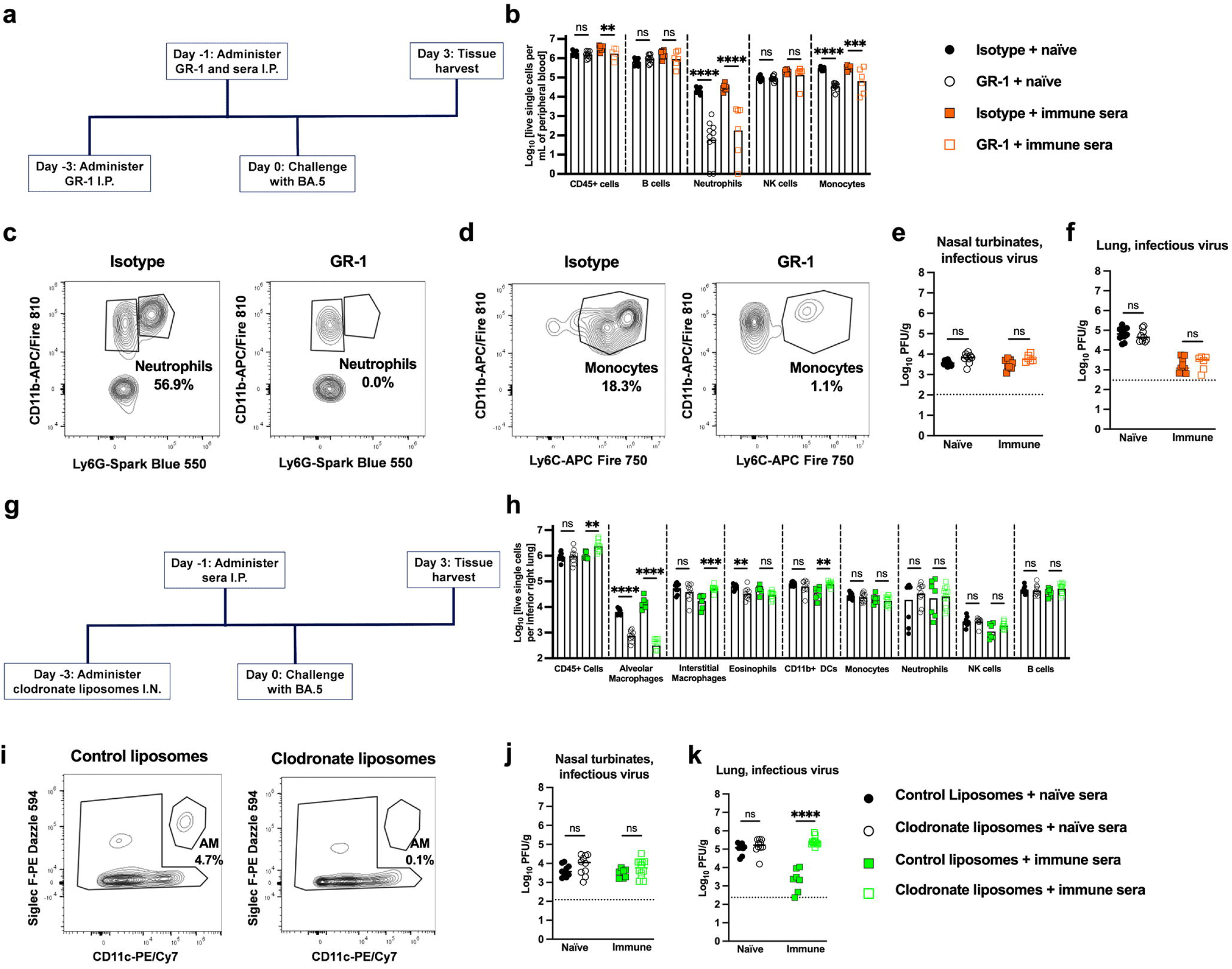
Depletion of alveolar macrophages impairs the protective activity of passively-transferred immune sera against BA.5 infection. (**a**) Scheme for depletion of neutrophils and monocytes, passive transfer of immune sera, and BA.5 challenge. Wild-type C57BL/6 twelve-week-old male mice were administered 500 μg of anti-Gr-1 (Ly6C/Ly6G) or isotype control antibody at Day −3 and −1 by intraperitoneal injection. On Day −1, mice were also given 60 μL of naïve or immune sera by intraperitoneal injection. On Day 0, mice were inoculated with 10^4^ FFU of BA.5, and tissues were harvested for virological analysis on Day +3. (**b-d**) Analysis of depletion of immune cell subsets in blood of mice receiving anti-Gr-1 (Ly6C/Ly6G) or isotype antibody at 3 dpi. Summary of different cell types (**b**) based on the gating strategy (see **Extended Data Fig 5a**). Results are from two experiments (n = 6-10 mice per group; Mann-Whitney test with Bonferroni post-test correction, ns, not significant; \*\**P* < 0.01, \*\*\**P* < 0.001\*\*\*\**P* < 0.0001). Representative flow cytometry dot plots showing depletion of neutrophils (**c**) and monocytes (**d**) with numbers indicating the cell population as a percentage of CD45^+^ cells. (**e-f**) Analysis of infectious viral burden by plaque assay at 3 dpi in nasal turbinates (**e**) and lungs (**f**) after BA.5 challenge (two experiments, n = 6-10 mice per group; Mann-Whitney test, ns, not significant). (**g**) Scheme for depletion of alveolar macrophages, passive transfer of immune sera, and BA.5 challenge. Wild-type C57BL/6 twelve-week-old male mice were administered control or clodronate liposomes at Day −3 by an intranasal route. On Day −1, mice were given 60 μL of naïve or immune sera by intraperitoneal injection. On Day 0, mice were inoculated with 10^4^ FFU of BA.5, and tissues were harvested on Day +3. (**h**) Analysis of depletion of immune cell subsets in lungs of mice receiving liposomes at 3 dpi. Summary of different cell types (**h**) based on gating strategy (see **Extended Data Fig 5b**). Results are from two experiments (n = 7-12 mice per group; Mann-Whitney test with Bonferroni post-test correction, ns, not significant; \*\**P* < 0.01, \*\*\**P* < 0.001, \*\*\*\**P* < 0.0001). (**i**) Representative flow cytometry dot plots showing depletion of alveolar macrophages after liposome administration with numbers indicating the cell population as a percentage of CD45^+^ cells. (**j-k**) Analysis of infectious viral burden by plaque assay at 3 dpi in nasal turbinates (**j**) and lungs (**k**) after BA.5 challenge (two experiments, n = 7-12 mice per group; Mann-Whitney test, ns, not significant; \*\*\*\**P* < 0.0001).

## DISCUSSION

Despite the diminished neutralizing ability of vaccine-elicited antibodies against SARS-CoV-2 variants with amino acid substitutions in the RBD and NTD^12,21^, protection against severe disease is maintained in the majority of vaccine recipients^36,37^. Although neutralizing activity of antibodies is a correlate of vaccine-mediated protection^4^, the ability of monovalent COVID-19 vaccines to protect against Omicron disease in the setting of waning serum antibody neutralization suggests additional protective immune mechanisms. These include anamnestic B cell responses that rapidly generate cross-reactive neutralizing antibodies, cross-reactive T cells responses, and/or non-neutralizing, cross-reactive antibodies that promote Fc mediated effector activities^6,16,18,38,39^. In our experiments, we focused on evaluating Fc mediated effector functions as a possible mechanism of vaccine-mediated protection against antigenic variants. *In vitro* studies with human convalescent sera have demonstrated that Fc effector functions are retained against antigenically variant strains, and that sera of COVID-19 patients with more severe disease have compromised FcγR binding abilities and effector functions^6,15,19,40^ Studies in mice with passively transferred mAbs show that Fc effector functions contribute to protection^41,42^, and this activity is maintained against antigenically distant strains even when neutralizing capacity is compromised^20^. Here, our experiments in mice show that Fc-FcγR interactions contribute to control of SARS-CoV-2 infection *in vivo* in the context of active or passive immunization, and that alveolar macrophages are a key cell type required for this activity.

Pooled immune serum from vaccinated mice was profiled for anti-spike antibodies against the ancestral SARS-CoV-2 strain and several variants of concern. The increased binding of immune antibodies to spike proteins was associated with several Fc effector functions including ADCP, ADNP, and ADCD. When immune serum was passively transferred to mice, protection against infection by SARS-CoV-2 strains MA-10 or WA1/2020 N501Y/D614G in the lungs required FcγR expression, particularly FcγR III, even though serum antibody neutralizing activity was present after transfer. These data showing a requirement for Fc effector functions for optimal antibody-mediated control of virus infection in the lung are consistent with studies in mice and hamsters with neutralizing mAbs that bind epitopes in the RBD^20,41^. Moreover, Fc-engineered anti-SARS-CoV-2 non-neutralizing and neutralizing mAbs binding the NTD and RBD regions confer greater protection in mice and hamsters^43,44^. In the context of passive antibody transfer, we observed less impact in the upper respiratory tract tissues, which could reflect the diminished ability of IgG antibodies in sera to accumulate in airway spaces^26^. Nonetheless, we observed FcγR-dependent reductions in viral load in the nasal turbinates after active mRNA vaccination, which could be due to higher levels of systemic anti-spike IgG/IgA or production of antibody by tissue-resident B cells. Persistent SARS-CoV-2 IgG antibodies in oral mucosal fluid and upper respiratory tract specimens have been reported following mRNA vaccination^45^.

In wild-type and FcγR KO mice immunized with mRNA-1273, we observed similar levels of neutralizing and RBD or spike-specific antibodies, and vaccine-induced CD4^+^ and CD8^+^ T cell responses. Although these results contrast with the idea that FcγRs have key roles in regulating adaptive immunity^46,47^, they are consistent with studies showing a lack of impairment of adaptive immune responses in FcγR KO mice to bacterial infection or IgG complexes^48^. Indeed, in control mRNA vaccinated animals, SARS-CoV-2 viral loads were similar in wild-type and FcγR KO mice. Thus, we attribute the diminished control of infection of the antigenically-shifted BA.5 strain in the turbinates and lungs of mRNA-1273 vaccinated FcγR KO mice to the loss of Fc-FcγR interactions that mediate antibody effector functions. Notwithstanding these results, antigen-matched bivalent mRNA vaccines targeting BA.1 and BA.5 spike proteins can induce higher levels of neutralizing antibodies against Omicron strains^8,33,49^, which might result in less reliance on Fc effector functions for protection against infection. We also performed cellular depletions to investigate the cell type responsible for the protection conferred by passively transferred antibody. Although several myeloid cells (monocytes, neutrophils, interstitial macrophages, and alveolar macrophages) in the lung express multiple FcγRs including FcγR III, depletion of alveolar macrophages compromised protection against infection. This effect was antibody-dependent since in the absence of immune sera, clodronate-dependent depletion of alveolar macrophages did not affect the viral burden in the lungs. These results are consistent with studies of influenza virus in mice, which showed that protective immunity conferred by non-neutralizing antibodies required alveolar macrophages and other lung phagocytes^35^.

### Limitations of study

We note several limitations in our study. (i) The conclusions showing an antiviral role for Fc effector functions of antibodies and FcγR III are based on experiments in mice. We used mice because of the availability of animals deficient in specific FcγRs, the reagents to achieve immune cell depletions, and the ability to perform both passive and active immunization, and BA.5 challenge. However, our results may not directly correlate with results in humans because of species-dependent differences in FcγR subtypes, functions, and expression on specific immune cells in the lung (**Supplementary Table 1**)^50,51^. Future studies with human FcγR transgenic mice^52^ lacking individual human FcγRs may help to bridge this gap. (ii) While we evaluated anti-spike antibody function in FcγR I KO, FcγR II KO, FcγR III KO, and FcγR I/III/IV KO mice, we did not directly assess a role for FcγR IV, as we did not have these KO mice. Passive and active immunization studies in FcγR IV KO mice^53^ are warranted. (iii) Although our studies in mice indicate an importance of alveolar macrophages and FcγR III interactions with antibody, we did not identify a specific cellular mechanism of action. While our serum profiling analysis is consistent with an antibody- and Fc-dependent phagocytic mechanism, it remains unclear if this occurs with opsonized virus or infected cells. (iv) We challenged mice with an antigenically-shifted BA.5 isolate. Future infection experiments using BQ.1.1, BF.7, or other emerging strains may be informative for determining the breadth of this mechanism of protection. (v) Although challenge of wild-type and FcγR KO C57BL/6 mice with BA.5 allowed us to evaluate the effects on viral burden in the setting of low levels of transferred or induced serum antibody neutralization, clinical disease and pathology do not develop because Omicron strains are less virulent in C57BL/6 and other strains of mice^54^. Testing of vaccinated wild-type and FcγR KO mice with antigenically-shifted yet more pathogenic SARS-CoV-2 strains [for mice] might enable assessment of the contribution of polyclonal antibodies and Fc effector functions to protection against lung inflammation. It will be important to determine whether vaccine-elicited antibodies engage FcγRs (*e.g*., FcγR I and FcγR III) on specific myeloid cells and promote inflammation, as infection-induced antibodies from patients enhanced SARS-CoV-2 uptake by monocytes and macrophages, and triggered inflammasome activation, pyroptotic cell death, and COVID-19 pathogenesis^55,56^. However, these studies also showed that immune plasma from mRNA vaccine recipients did not promote antibody-dependent monocyte infection and inflammation.

In summary, our experiments in mice provide insight and help to explain human studies that correlate Fc-FcγR interactions with clinical outcome against SARS-CoV-2 and emerging variants of concern. We demonstrate the importance of particular murine FcγRs in mediating antibody protection against SARS-CoV-2 infection and identify alveolar macrophages as a key contributing cell type in the context of passive immunization. Our results also provide an explanation as to how Fc-FcγR interactions might contribute to monovalent vaccine-mediated protection against severe infection by SARS-CoV-2 variants even in the setting when serum neutralizing antibody activity is lost^7^. They also suggest that targeting Fc effector functions in the context of vaccine design could be a strategy to generate more broadly protective immune responses^23^. Future studies are warranted to define the epitopes targeted by antibodies with strong Fc effector functions and develop improved *in vitro* Fc effector function assays that correlate better with protection *in vivo* against infection by SARS-CoV-2 and variants^57^.

## Acknowledgements

This study was supported by the NIH (R01 AI157155, NIAID Centers of Excellence for Influenza Research and Response (CEIRR) contracts 75N93021C00014 and 75N93019C00051, to M.S.D.; and R01 AI110700 to R.S.B.). We thank Mehul Suthar for the BA.5 isolate used in this study and Taia Wang for critical comments on the manuscript.

## Author contributions

S.R.M. performed binding and neutralization assays, immunization, passive transfer, depletion studies, challenge experiments, and flow cytometry. P.D. performed and analyzed T cell responses. B.W. performed mouse experiments. C.E.K. performed some of the flow cytometry experiments. M.L. performed immune cell processing and staining. R.S.B. provided the mouse-adapted MA-10 strain. R.P.M., T.M.C., and G.A. designed, performed, and analyzed the serological profiling experiments. D.K.E. provided mRNA vaccines and helped to design vaccination experiments. S.R.M. and M.S.D. designed studies and wrote the initial draft, with the other authors providing editorial comments.

## EXTENDED DATA FIGURES

**Extended Data Figure 1. Gating strategy for Luminex-based and Fc effector function assays.** (**a**) Gating for Luminex-bead based antibody binding to spike-coated beads. (**b**) Gating for ADNP assay showing CD66^+^ neutrophils with opsinophagocytosed beads. (**c**) Gating for ADCP assay showing THP-1 monocytes and opsinophagocytosed beads. (**d**) Gating for ADCD assay showing complement deposition on spike and antibody coated beads. (**e**) Gating for NK cell activation assay showing CD107a expression.

**Extended Data Figure 2. T cell responses in mRNA-1273 vaccinated wild-type and FcγR KO mice.** (**a**) Representative flow cytometry plots show gating scheme for quantification of spike-specific CD4^+^ and CD8^+^ T cell responses in the spleen of wild-type, FcγR III KO, and FcγR I/III/IV KO mice at day 10 after boosting with control or mRNA-1273 vaccines. (**b-c**) At day 10 after boosting, the spleen of wild-type and FcγR KO mice were harvested, and T cell responses were measured *ex vivo* after spike peptide re-stimulation. Splenocytes were incubated overnight with class I MHC (**b**) or class II MHC (**c**) immunodominant spike peptides, and the percentages and numbers of IFNγ and TNFα positive CD8^+^ (**b**) or CD4^+^ (**c**) T cells were quantified by intracellular staining and flow cytometry. Data are pooled from two experiments (n = 9-10 per group). Comparisons were made between groups that received the mRNA 1273 vaccine (one-way ANOVA with Tukey’s post-test; all comparisons were not significant; column height indicates mean values).

**Extended Data Figure 3. Levels of anti-BA.5 antibody in mice passively transferred vaccine-elicited serum antibody.** (**a**) Levels of anti BA.5 spike IgG in serum of mice that were passively transferred naïve or immune sera. Amounts are compared to those in vaccine-elicited immune serum before transfer (n = 5 mice per group, columns indicate mean values). (**b**) Neutralizing antibody response against SARS-CoV-2 BA.5 using sera from naïve (circles) or spike protein vaccinated (grey squares) mice. Also shown is serum neutralizing antibody activity from recipient wild-type mice one day after transfer of immune sera (black squares). The data are representative of results with n = 5 mice per group.

**Extended Data Figure 4. FcγR expression on myeloid cells in the lung.** Lung cells from wild-type (black), FcγR I KO (purple), FcγR III KO (blue), and FcγR I/III/IV KO (green) mice were strained with antibodies for FcγR I, FcγR III, or FcγR IV. After gating on live cells, alveolar macrophages, neutrophils, and monocytes were defined (see **Extended Data Fig 5**). The data are representative of results with n = 3 mice per group, and histograms are shown.

**Extended Data Figure 5. Gating scheme for analysis of cell populations in the blood and lung.** (**a**) Immune cell populations in the blood of C57BL/6 mice were analyzed using the indicated gating scheme and conjugated antibodies. After gating on live single cells, monocytes (P5) were defined as CD45^+^ CD11b^hi^ Ly6C^hi^; neutrophils (P6) were defined as CD45^+^ CD11b^hi^ Ly6G^hi^; natural killer (NK) cells (P7) were defined as CD45^+^ NK1.1^+^; and B cells (P8) were defined as CD45^+^ B220^+^ cells. (**b**) Immune cell populations in the lungs of C57BL/6 mice were analyzed using the indicated gating scheme and conjugated antibodies. After gating on live single cells, alveolar macrophages (P1) were defined as CD45^+^ CD11c^+^ Siglec-F^+^; interstitial macrophages (P2) were defined as CD45^+^ CD64^+^, eosinophils (P3) were defined as CD45^+^ CD11b^+^ Siglec-F^+^; CD11b dendritic cells (P4) were defined CD45^+^ CD11b^+^ CD11c^+^, Siglec-F^-^, MHC II^+^; monocytes (P5) were defined as CD45^+^ Ly6C^hi^; neutrophils (P6) were identified as CD45^+^ Ly6G^+^; natural killer (NK) cells (P7) were defined as CD45^+^ NK1.1^+^; and B cells (P8) were defined as CD45^+^ B220^+^.

---

**Supplementary Table 1.** Murine and human FcgR expression in immune cells of the lung

| Lung cell subset | Human FcγRs |  |  |  |  |  |
| --- | --- | --- | --- | --- | --- | --- |
|  | FcγR I<br>(CD64)<br>activating | FcγR IIa<br>(CD32a)<br>activating | FcγR IIb<br>(CD32b)<br>inhibitory | FcγR IIc<br>(CD32c)<br>activating | FcγR IIIa<br>(CD16a)<br>activating | FcγR IIIb<br>(CD16b)<br>(GPI-anchored) |
| Alveolar macrophages | + | + | + | - | + | - |
| Interstitial macrophages | + | + | + | - | + | - |
| Ly6C<sup>+</sup> monocytes | + | + | + | - | - | - |
| Neutrophils | Inducible | + | + | - | - | + |
| Eosinophils | Inducible | + | + | - | - | Inducible |
| CD11b<sup>+</sup> Dendritic cells | + | + | + | - | Inducible | - |
| NK cells | - | - | - | + | + | - |
| B cells | - | - | + | - | - | - |

| Lung cell subset | Mouse FcγRs |  |  |  |
| --- | --- | --- | --- | --- |
|  | FcγR I<br>(CD64)<br>activating | FcγR IIb<br>(CD32b)<br>inhibitory | FcγR III<br>(CD16)<br>activating | FcγR IV<br>(CD16-2)<br>activating |
| Alveolar macrophages | + | + | + | + |
| Interstitial macrophages | + | + | + | + |
| Ly6C<sup>+</sup> monocytes |  | + | + | + |
| Neutrophils | - | + | + | + |
| Eosinophils | - | + | Inducible | - |
| CD11b<sup>+</sup> Dendritic cells | + | + | + | - |
| NK cells | - | - | + | - |
| B cells | - | + | - | - |

Table was compiled based on published results<sup>51,58-60</sup>.

## METHODS

### Cells

Vero-TMPRSS2^61^ and Vero-hACE2-TMPRSS2^62^ cells were cultured at 37°C in Dulbecco’s Modified Eagle medium (DMEM) supplemented with 10% fetal bovine serum (FBS), 10□mM HEPES pH 7.3, 1□mM sodium pyruvate, 1× non-essential amino acids, and 100□U/mL of penicillin–streptomycin. Vero-TMPRSS2 and Vero-hACE2-TMPRSS2 cells were supplemented with 5 μg/mL of blasticidin and 10 μg/mL of puromycin, respectively. All cells routinely tested negative for mycoplasma using a PCR-based assay.

### Viruses

All SARS-CoV-2 strains used (WA1/2020 N501Y/D614G, mouse-adapted MA-10, B.1.351, and BA.5) have been previously described^25,33,62,63^. All viruses were subjected to next generation deep sequencing to confirm presence and stability of substitutions. All experiments with virus were performed in approved biosafety level 3 (BSL-3) facilities with appropriate positive-pressure respirators, personal protective gear, and containment.

### Mice

Animal studies were carried out in accordance with the recommendations in the Guide for the Care and Use of Laboratory Animals of the National Institutes of Health. The protocols were approved by the Institutional Animal Care and Use Committee at the Washington University School of Medicine (assurance number A3381–01). Virus inoculations were performed under anesthesia that was induced and maintained with ketamine hydrochloride and xylazine, and all efforts were made to minimize animal suffering. Experiments were neither randomized nor blinded. C57BL/6J mice (Cat # 000664) were obtained from The Jackson Laboratory. FcγR I KO^64^, FcγR II KO (Taconic Biosciences; Cat # 580), FcγR III KO (Jackson Laboratory; Cat # 009637), FcγR I/III/IV (common γ-chain) KO (Taconic Biosciences; Cat # 583), and C1q KO^65^ were obtained commercially or collaborators and then backcrossed onto a C57BL/6J background (>99%) using Speed Congenics (Charles River Laboratories) and single nucleotide polymorphism analysis. Animals were housed in groups and fed standard chow diets.

### Preclinical mRNA and ChAd-SARS-CoV-2 vaccines

A sequence-optimized mRNA encoding prefusion-stabilized Wuhan-Hu-1 (mRNA-1273) was designed and synthesized *in vitro* using an optimized T7 RNA polymerase-mediated transcription reaction with complete replacement of uridine by N1m-pseudouridine^66^. A non-translating control mRNA was synthesized and formulated into lipid nanoparticles as previously described^67^. The reaction included a DNA template containing the immunogen open-reading frame flanked by 5’ untranslated region (UTR) and 3’ UTR sequences and was terminated by an encoded polyA tail. After RNA transcription, the cap-1 structure was added using the vaccinia virus capping enzyme and 2 -*O*-methyltransferase (New England Biolabs). The mRNA was purified by oligo-dT affinity purification, buffer exchanged by tangential flow filtration into sodium acetate, pH 5.0, sterile filtered, and kept frozen at −20°C until further use. The mRNA was encapsulated in a lipid nanoparticle through a modified ethanol-drop nanoprecipitation process described previously^68^. Ionizable, structural, helper, and polyethylene glycol lipids were briefly mixed with mRNA in an acetate buffer, pH 5.0, at a ratio of 2.5:1 (lipid:mRNA). The mixture was neutralized with Tris-HCl, pH 7.5, sucrose was added as a cryoprotectant, and the final solution was sterile-filtered. Vials were filled with formulated lipid nanoparticle and stored frozen at −20°C until further use.

The ChAd-SARS-CoV-2 and ChAd-Control vaccine vectors were derived from simian Ad36 backbones^69^, and the constructing and validation has been described in detail previously^70^. The rescued replication-incompetent ChAd-SARS-CoV-2-S were scaled up in HEK293 cells and purified by CsCl density-gradient ultracentrifugation. For passive transfer studies, a large batch (8 mL) of immune sera (obtained ≥ 30 days post immunization) was pooled from C57BL/6 mice vaccinated with ChAd-SARS-CoV-2 or mRNA-1273.

### Viral antigens

Recombinant RBD proteins from Wuhan-1 and BA.5 SARS-CoV-2 strains were expressed as described^71,72^. Recombinant proteins were produced in Expi293F cells (ThermoFisher) by transfection of DNA using the ExpiFectamine 293 Transfection Kit (ThermoFisher). Supernatants were harvested 3 days post-transfection, and recombinant proteins were purified using Ni-NTA agarose (ThermoFisher), then buffer exchanged into PBS and concentrated using Amicon Ultracel centrifugal filters (EMD Millipore).

### ELISA

Recombinant Wuhan-1, BA.4/5 receptor binding domain (RBD), or BA.4/5 spike protein (4 μg/mL) was immobilized on 96-well Maxisorp ELISA plates (Thermo Fisher) overnight at 4°C in coating buffer (1X PBS supplemented with 0.05% Tween-20, 2% BSA, and 0.02% NaN_3_). Plates were washed with PBS and blocked with 4% BSA for one hour at 25°C. Serum was serially diluted in 2% BSA and incubated on plates for 1 h at 25°C. After washing with PBS, RBD or spike-bound serum antibodies were detected with horseradish peroxidase conjugated goat anti-mouse IgG (1:500 dilution, Milipore Sigma) incubating for 2 h at 25°C. Plates were washed and developed with 3,3’-5,5’ tetramethylbenzidine substrate (Thermo Fisher), halted with 2 N H_2_SO_4_ and read at 450 nm using a microplate reader (BioTek). Mean serum endpoint titers were calculated with curve fit analysis of optical density (OD) values set as the reciprocal value of the serum dilution equal to the mean plus six times the standard deviation of background signal.

### Focus reduction neutralization test

Serial dilutions of sera were incubated with 10^2^ focus-forming units (FFU) of WA1/2020 N501Y/D614G or BA.5 for 1 h at 37°C. Antibody-virus complexes were added to Vero-TMPRSS2 cell monolayers in 96-well plates and incubated at 37°C for 1 h. Subsequently, cells were overlaid with 1% (w/v) methylcellulose in MEM. Plates were harvested 30 h (WA1/2020 N501Y/D614G and MA-10) or 72 h (BA.5) later by removing overlays and fixed with 4% PFA in PBS for 20 min at room temperature. Plates were washed and sequentially incubated with an oligoclonal pool (SARS2 −02, −08, −09, −10, −11, −13, −14, −17, −20, −26, −27, −28, −31, −38, −41, −42, −44, −49, −57, −62, −64, −65, −67, and −71^73^ of anti-S murine antibodies (including cross-reactive mAbs to SARS-CoV) and HRP-conjugated goat anti-mouse IgG (Sigma Cat # A8924, RRID: AB_258426) in PBS supplemented with 0.1% saponin and 0.1% bovine serum albumin. SARS-CoV-2-infected cell foci were visualized using TrueBlue peroxidase substrate (KPL) and quantitated on an ImmunoSpot microanalyzer (Cellular Technologies).

### Viral plaque assay

Titration of infectious SARS-CoV-2 was performed as previously described^74^. Briefly, lung and nasal turbinate homogenates were serially diluted and added to Vero-TMPRSS2-hACE2 cell monolayers in 24-well tissue culture plates for 1 h at 37°C. Cells were then overlaid with 1% (w/v) methylcellulose in MEM and incubated for 72 h (MA-10 and WA1/2020 N501Y/D614G) or 96 h (BA.5). Subsequently, cells were fixed with 4% paraformaldehyde in PBS for 20 min at room temperature before staining with 0.05% (w/v) crystal violet in 20% methanol. Viral plaques were counted manually.

### Mouse experiments

(a) Passive transfer studies. Twelve-week-old male C57BL/6, FcγR I KO, FcγR II KO, FcγR III KO, FcγR I/III/IV KO, and C1q KO mice were administered 60 ul of sera (naïve or pooled from mice immunized and boosted with mRNA-1273 or ChAd-SARS-CoV-2-S) one day prior to challenge with 50 μl of 10^3^ FFU of MA-10 or 10^4^ FFU of WA1/2020 N501Y/D614G by intranasal administration. Lungs, nasal turbinates, and nasal washes (collected in 500 μl of .5% bovine serum albumin in phosphate buffered saline) were harvested four days after inoculation for virological analysis. (b) Immunization studies. Nine-week-old male C57BL/6, FcγR I KO, FcγR III KO, and FcγR I/III/IV KO mice were immunized and boosted with 0.25 μg of control or mRNA-1273 vaccine by intramuscular injection four weeks apart. Animals were bled twenty-four days after boosting for immunogenicity analysis of sera and then challenged four days later with 50 μl of 10^4^ FFU of BA.5 by intranasal administration. Lungs, nasal turbinates, and nasal washes (collected in 500 μl of .5% bovine serum albumin in phosphate buffered saline) were harvested three days after inoculation for virological analysis. In vivo studies were not blinded with mice randomly assigned to treatment groups.

### Immune cell depletions

For monocyte and neutrophil depletions, anti-Ly6G/Ly6C (BioXCell; clone RB8-8C5; 500 μg) or an IgG2b isotype control (BioXCell; clone LTF-2; 500 μg) were administered to mice by intraperitoneal injection at days −3 and −1 relative to SARS-CoV-2 inoculation. For alveolar macrophage depletion, clodronate liposomes (Liposoma; 250 μg) or control liposomes (Liposoma; 250 μg) were administered via intranasal route at day −3 relative to SARS-CoV-2 inoculation.

For analysis of neutrophil and monocyte depletion, peripheral blood was collected on the day of harvest. Erythrocytes were lysed with ACK lysis buffer (GIBCO) at room temperature for 3 min and resuspended in RPMI 1640. Single-cell suspensions were preincubated with Fc block antibody (1:100; clone S17011E; Biolegend) in PBS + 2% heat-inactivated FBS + 1 mM EDTA for 20 min at 4°C, stained with antibodies against CD45 (AF488; clone 30-F11; Biolegend), CD11b (APC/Fire 810; clone M1/70; Biolegend), Ly6G (Spark Blue 550; clone 1A8; Biolegend), Ly6C (APC-Fire 750; clone HK1.4; Biolegend), NK1.1 (BV570; clone PK136; Biolegend), B220 (Pacific Blue; clone RA3-6B2; Biolegend), fixable viability dye (eFluor 506; BD Biosciences), True-Stain Monocyte Blocker (Biolegend; 5 μl/sample), and Brilliant Stain Buffer Plus (BD Biosciences; 10 μl/sample), and then fixed with 4% paraformaldehyde in PBS for 20 min at room temperature. All antibodies were used at a dilution of 1:200. The viability dye was used at 1:100. Absolute cell counts were determined using Precision Count Beads (Biolegend). Flow cytometry data were acquired on a 3 laser Cytek Aurora cytometer (Cytekbio) and analyzed using FlowJo software v10.8 (Treestar).

For analysis of lung tissues, mice were euthanized by ketamine overdose, perfused with sterile PBS, and the right inferior lung lobes were digested in 50 μL of 5 mg/mL of Liberase Tm (Roche) and 12.5 μL of 10 mg/mL of DNase I (Sigma-Aldrich) in 5 mL of HBSS at 37°C for 35 min. Single cell suspensions of lung digest were preincubated with Fc block antibody (Biolegend) in PBS + 2% heat-inactivated FBS + 1 mM EDTA for 20 min at 4°C. Cells were then stained with antibodies against CD45 (AF488), CD11b (APC/Fire 810), MHC II (BV711; clone M5/114.15.2; Biolegend), CD11c (PE-Cy7; clone N418; Biolegend), CD64 (BV421; clone X54-5/7.1; Biolegend), CD88 (Viogreen; clone REA1206; Miltenyi Biotec), Siglec-F (PE Dazzle 594; clone S17007L; Biolegend), Ly6G (Spark Blue 550), Ly6C (APC-Fire 750), NK1.1 (BV570), CD3 (BV650; clone 145-2C11; BD Biosciences), B220 (Pacific Blue), CD16.2 (APC), CD16 (PE), fixable viability dye (eFluor 506), True-Stain Monocyte Blocker (5 μl/sample), and Brilliant Stain Buffer Plus (10 μl/sample) and then fixed with 4% paraformaldehyde in PBS for 20 min at room temperature. All antibodies were used at a dilution of 1:200 with the exceptions of MHC II BV711, which was used at 1:300 and the viability dye, used at 1:100. In experiments staining for mouse FcγR III, Fc block was not used. Absolute cell counts were determined using Precision Count Beads (Biolegend). Flow cytometry data were acquired on a 3 laser Cytek Aurora cytometer (Cytekbio) and analyzed using FlowJo software (version 10.4.2).

### Measurement of viral burden

Tissues were weighed and homogenized with zirconia beads in a MagNA Lyser instrument (Roche Life Science) in 1 ml of DMEM medium supplemented with 2% heat-inactivated FBS. Tissue homogenates were clarified by centrifugation at 10,000 rpm for 5 min and stored at −80°C. RNA was extracted using the MagMax mirVana Total RNA isolation kit (Thermo Fisher Scientific) on the Kingfisher Flex extraction robot (Thermo Fisher Scientific). RNA was reverse transcribed and amplified using the TaqMan RNA-to-CT 1-Step Kit (Thermo Fisher Scientific). Reverse transcription was carried out at 48°C for 15 min followed by 2 min at 95°C. Amplification was accomplished over 50 cycles as follows: 95°C for 15 s and 60°C for 1 min. Copies of SARS-CoV-2 *N* gene RNA in samples were determined using a published assay^74^.

### Antibody isotype and Fc-receptor binding profiling

Serum samples were analyzed by a customized Luminex assay to quantify the levels of antigen-specific antibody subclasses and FcγR binding profiles, as previously described^75,76^. Briefly, antigens were coupled to magnetic Luminex beads (Luminex Corp) by carbodiimide-NHS ester-coupling (Thermo Fisher). The antigen-coupled microspheres were washed and incubated with heat-inactivated serum samples at an appropriate sample dilution (1:100-1:400 for antibody isotyping and 1:1,000 for all low-affinity FcγRs) overnight in 384-well plates with continuous shaking (Greiner Bio-One). Unbound antibodies were washed away using the magnetic 384-well HydroSpeed Plate Washer (Tecan) using 1X Luminex assay buffer (1X PBS, 0.1% BSA, 0.05% Tween-20). Secondary antibodies (PE-coupled IgG1, IgG2b, IgG2c, IgG3) were added and incubated for 1 h at room temperature with continuous shaking. Unbound complexes were washed away using the magnetic 384-well Hydrospeed Plate Washer. Beads were resuspended in 40 μL of QSOL buffer (Sartorius) and then run on the iQue Screener PLUS (Intellicyt) using a customized gating strategy for each bead region (**Extended Data Fig 1**). Median fluorescence intensity was calculated for all samples, which were run in technical replicates.

For FcγR binding, sera were incubated with antigen-coated beads and washed as described above. Custom synthesized FcγR (FcγR2b, FcγR3, FcγR4; Duke Protein Production facility) were biotinylated and then bound to PE-Streptavidin. The labeled FcγRs were then incubated with the sera for 1 h at room temperature with continuous shaking. Unbound complexes were washed as indicated above. Beads were resuspended in 40 μL of QSOL buffer and then run on the iQue Screen PLUS (Intellicyt). All flow cytometry files were analyzed using Intellicyt ForeCyt (v8.1).

All antigens and FcγRs were equilibrated in 1X PBS using Zeba-Spin desalting and size exclusion chromatography columns (ThermoFisher) prior to bead coupling. Dilution curves for each antibody isotype and subclass and FcγRs were performed for each antigen to ensure reported values were within the linear range of detection. Binding for antigens was calculated as the fold increase relative to naïve levels, which arbitrarily were set to 1.

### Evaluation of antibody-mediated effector functions

ADCP and ADNP were evaluated using a flow-cytometry-based phagocytic assay using fluorescently labeled microspheres. In brief, fluorescent neutravidin microspheres were coupled with biotinylated antigens and then incubated with the diluted serum to form immune complexes. Bead-bound immune complexes were incubated with monocytes (THP-1 cells) or neutrophils overnight at 37 °C in 96 well, round-bottom plates (Corning). After overnight incubation, plates were centrifuged and non-bound beads/immune complexes were removed. Cells were then fixed in 4% paraformaldehyde and stained for indicated markers. Microsphere-positive cells were quantified through flow cytometry and phagocytic scores were calculated as (the percentage of microsphere positive cells * MFI of positive cells) /100,000.

ADCD was quantified through the coupling of biotinylated antigens to neutravidin microspheres and then incubated with guinea pig complement at 37 °C for 50 min. Reactions were quenched with 15 mM EDTA. Fluorescein-conjugated anti-C3b was added to the mixture for 1 h with constant shaking. Plates were washed with 1X PBS and complexes were fixed with 4% paraformaldehyde and washed again with 1X PBS. Beads were resuspended in 35 μL of 1X PBS and then analyzed by flow cytometry. Complement deposition was calculated as the fold increase relative to naïve levels, which arbitrarily were set to 1.

ADNKA was quantified through the surface expression of CD107a (as a marker for degranulation). In brief, buffy coats were obtained from healthy donors at Massachusetts General Hospital and enriched using the RosetteSep NK enrichment kit (Stemcell) in the presence of IL-15. Antigen-coated, 96-well ELISA plates were then incubated with serum and the NK cell preparation. Plates were placed in a 37°C incubator for 2 h. Reactions were stopped and cells were fixed and stained extracellularly with anti-CD107a-PeCy5, anti-CD3-PB, anti-CD56-PE-Cy7, and anti-CD16-APC-Cy7. Cells were washed three times with 1X PBS and resuspended in 30 μL of 1X PBS and analyzed by flow cytometry.

### T cell analysis

Spleens were harvested from control or mRNA-1273 vaccinated wild type or FcγR KO mice at day 10 after boosting, and single cell suspensions were generated after tissue disruption and passage through a 70-μm cell strainer. Splenocytes were pelleted by centrifugation, and erythrocytes were lysed using ACK lysis buffer (Thermo Fisher). Cells then were re-suspended in RPMI 1640 media supplemented with 10% FBS, 1% HEPES, 1% L-glutamine and 0.1% β-mercaptoethanol. For peptide stimulation, splenocytes were incubated separately with class I MHC (VL8, peptide sequence S539-546: VNFNFNGL^32^) or class II MHC ((#62, peptide sequence S62-76: VTWFHAIHVSGTNGT^31^) immunodominant spike peptides (1 μg/mL) overnight at 37°C in the presence of Brefeldin A (1:500, Invitrogen). The following day, cells were washed and stained with Fc block (Clone 93; Cat: 101320; BioLegend), CD45 (BUV395; Clone 30-F11; Cat: 564279; BD Biosciences), CD8β (PerCP/Cy5.5; Clone YTS156.7.7; Cat: 126610; BioLegend), CD4 (FITC; Clone GK1.5; Cat: 100406; BioLegend), CD44 (APC/Cy7; Clone IM7; Cat: 103028; BioLegend) for 30 min at 4°C in FACS buffer (1x PBS with 2% FCS and 2 mM EDTA). Dead cells were excluded using Live/Dead (Thermo Fisher) that was added concurrently with staining. Following this, cells were washed, fixed with and stained for intracellular IFN-γ APC; Clone XMG1.2; Cat: 505810; BioLegend) and TNF-α (PE/Cy7; Clone MP6-XT22; Cat: 25-7321-82; Invitrogen) using BD fixation/permeabilization kit (BD Biosciences) according to the manufacturer’s instructions. Cells were processed by flow cytometry on a Cytek Aurora and analyzed using FlowJo software version 10.4.2).

### Statistical analysis

Statistical significance was assigned when *P* values were < 0.05 using GraphPad Prism version 9.3. Tests, number of animals, median values, and statistical comparison groups are indicated in the Figure legends. Changes in infectious virus titer, viral RNA levels, or serum antibody responses were analyzed by one-way ANOVA with a post-test correction when comparing three or more groups. When comparing two groups, a Mann-Whiteny test was performed and a Bonferrroni correction was used to account multiple independent comparisons. Best-fit lines were calculated using non-linear regression analyses.

### Materials availability

All requests for resources and reagents should be directed to the Lead Contact author. This includes viruses, proteins, vaccines, and primer-probe sets. All reagents will be made available on request after completion of a Materials Transfer Agreement (MTA). The preclinical mRNA vaccines (control and mRNA-1273) can be obtained under an MTA with Moderna (contact: Darin Edwards).

### Data availability

All data supporting the findings of this study are available within the paper, its Extended Data, or Source Data files. Any additional information related to the study also is available from the corresponding author upon request.

### Code availability

No code was used in the course of the data acquisition or analysis.

