## Supplementary figures and images for "Fcγ receptor-dependent antibody effector functions are required for vaccine protection against infection by antigenic variants of SARS-CoV-2"

### Extended Data Figure 1

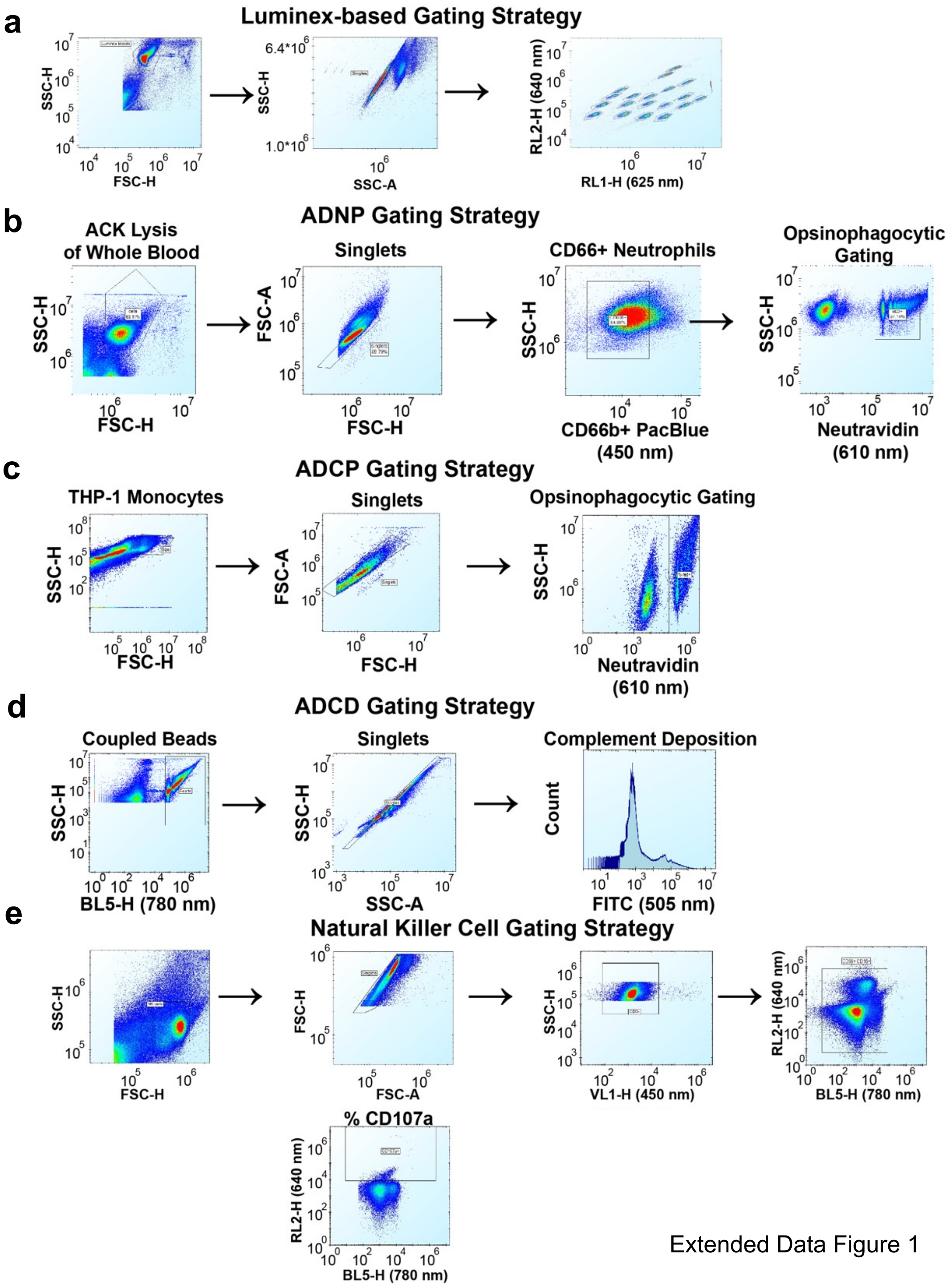


Extended Data Figure 1

### Extended Data Figure 2

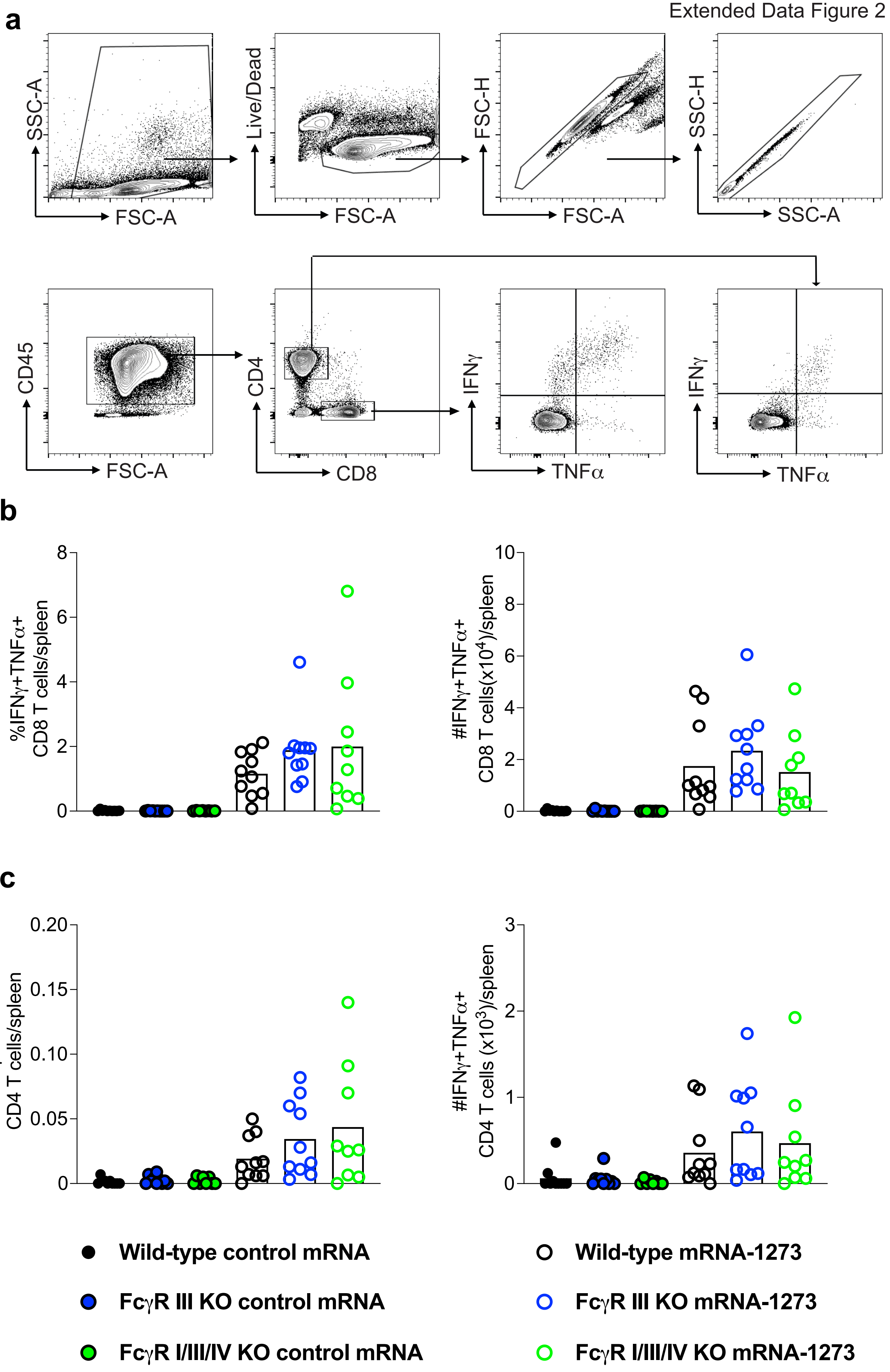

### Extended Data Figure 3

Extended Data Figure 3

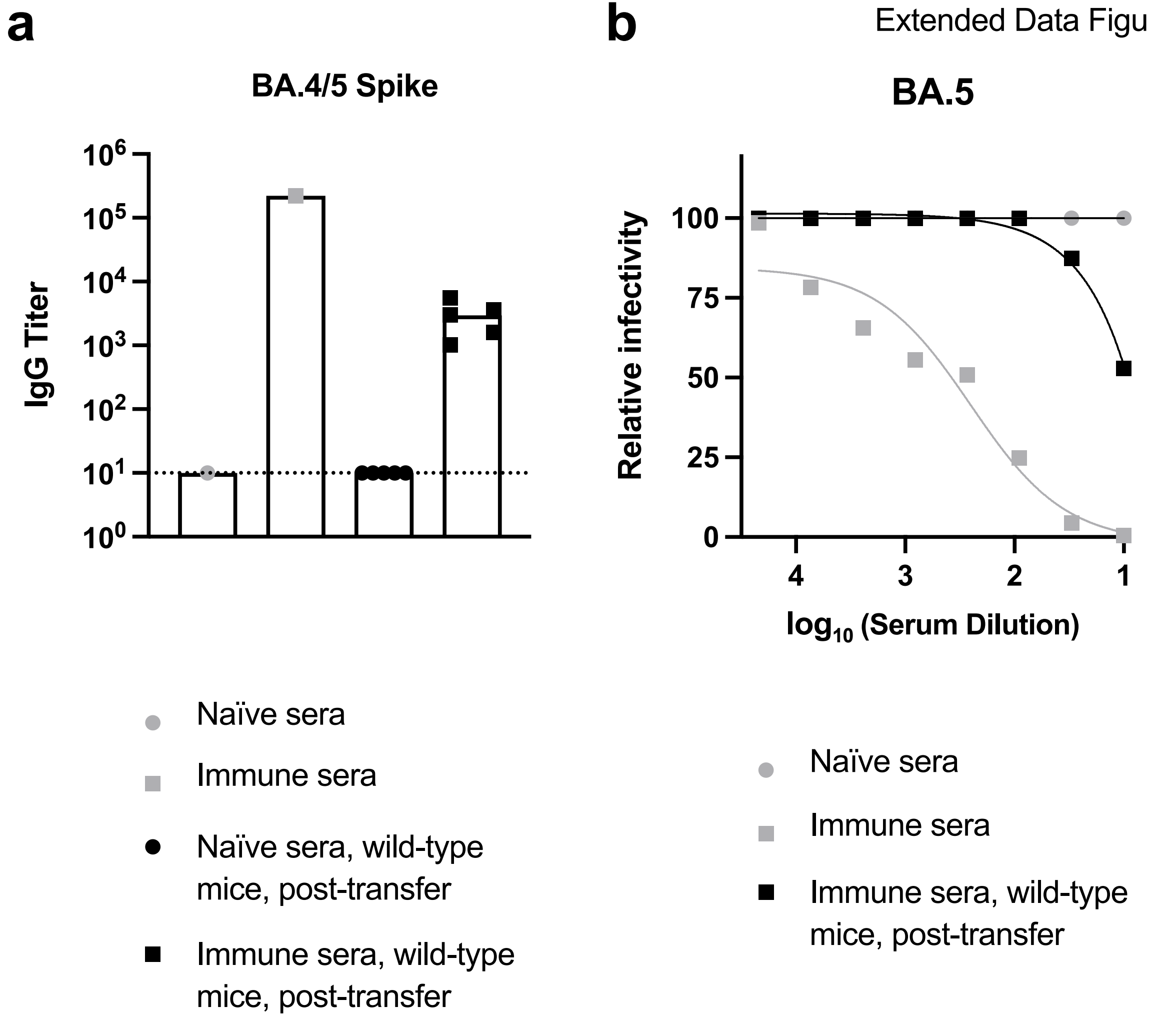

### Extended Data Figure 4

Extended Data Fig 4

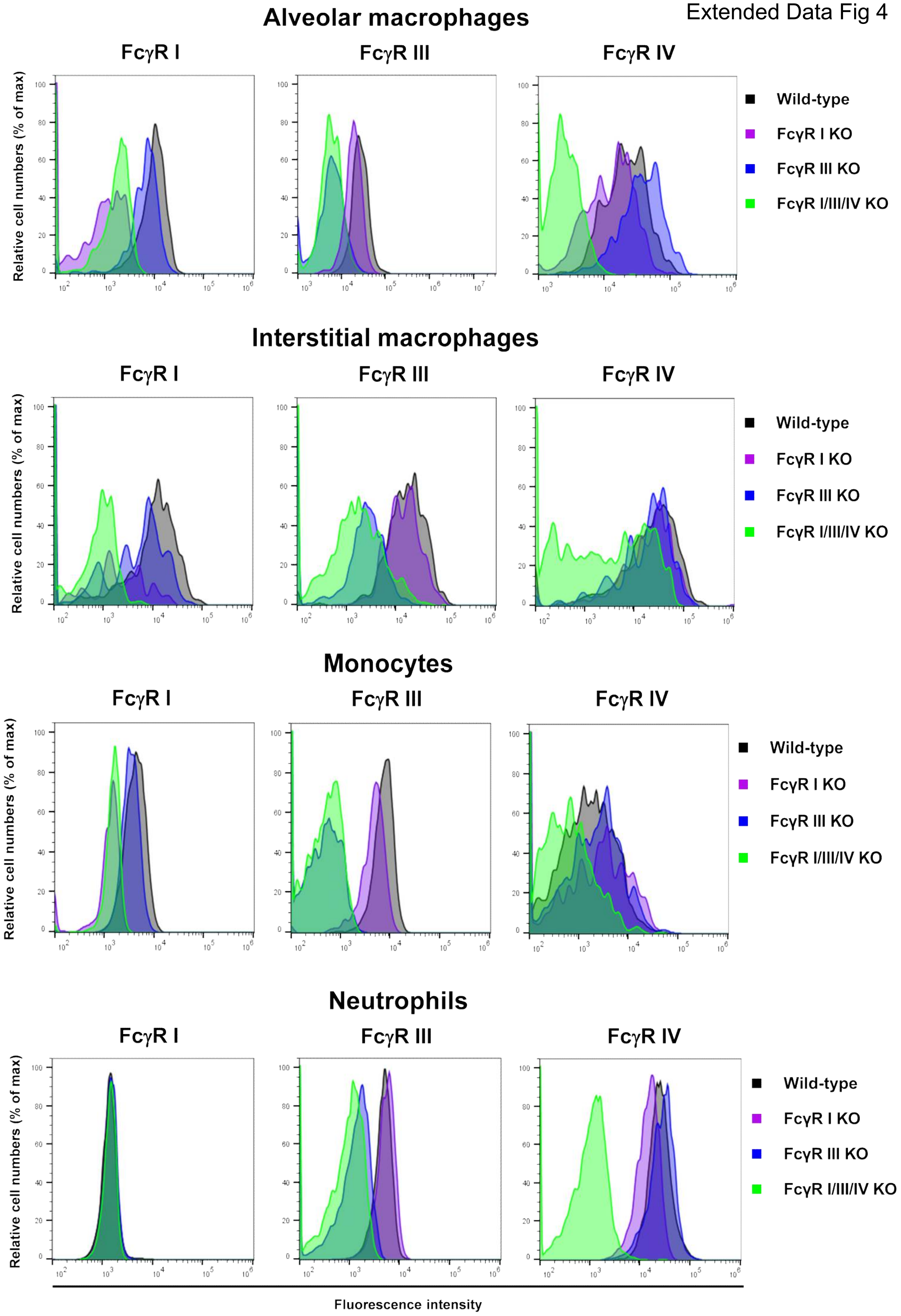
