## Extended Data Figure 4 for "Fcγ receptor-dependent antibody effector functions are required for vaccine protection against infection by antigenic variants of SARS-CoV-2"

a

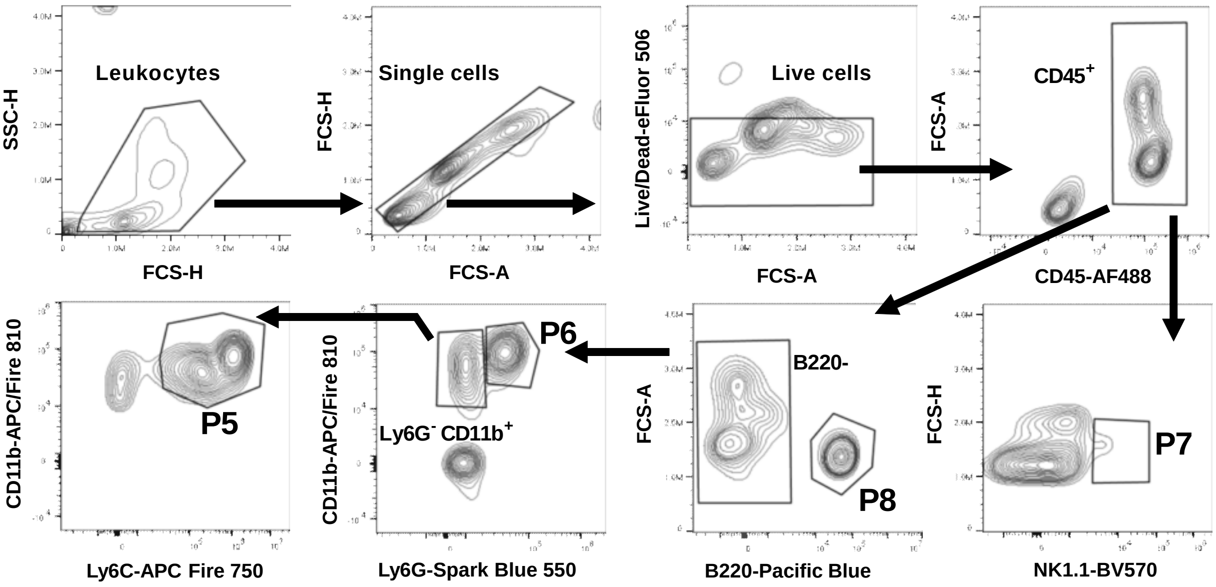


**P1: Alveolar macrophages**
**P2: Interstitial macrophages**
**P3: Eosinophils**
**P4: CD11b$^{+}$ DCs**

**P5: Monocytes**
**P6: Neutrophils**
**P7: NK cells**
**P8: B cells**

b

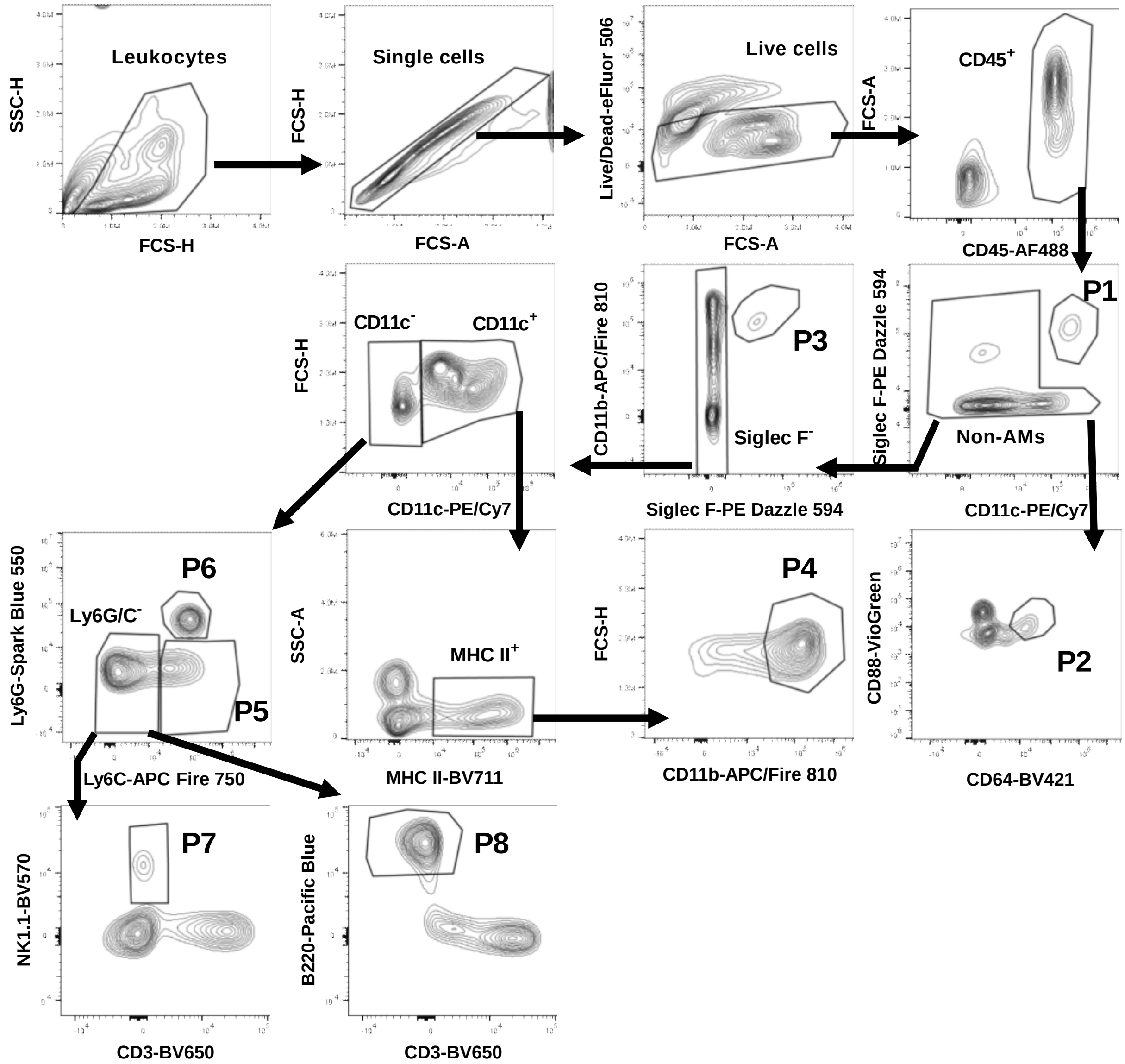
